# The Sle1 Cell Wall Amidase Controls Daughter Cell Splitting, Cell Size, and β-Lactam Resistance in Community Acquired Methicillin Resistant *Staphylococcus aureus* USA300

**DOI:** 10.1101/772616

**Authors:** Ida Thalsø-Madsen, Fernando Ruiz Torrubia, Lijuan Xu, Andreas Petersen, Camilla Jensen, Dorte Frees

## Abstract

Most clinically relevant methicillin resistant *Staphylococcus aureus* (MRSA) strains have become resistant to β-lactams antibiotics through horizontal acquisition of the *mecA* gene encoding PBP2a, a peptidoglycan transpeptidase with low affinity for β-lactams. The level of resistance conferred by *mecA* is, however, strain dependent and the mechanisms underlying this phenomenon remain poorly understood. We here show that β-lactam resistance correlates to expression of the Sle1 cell wall amidase in the fast spreading and highly virulent community-acquired MRSA USA300 clone. Sle1 is a substrate of the ClpXP protease, and while the high Sle1 levels in cells lacking ClpXP activity confer β-lactam hyper-resistance, USA300 cells lacking Sle1 are as sensitive to β-lactams as cells lacking *mecA*. This finding prompted us to assess the cellular roles of Sle1 in more detail, and we demonstrate that high Sle1 levels accelerate the onset of daughter cells splitting and decrease cell size. Vice versa, oxacillin decreases the Sle1 level, and imposes a cell-separation defect that is antagonized by high Sle1 levels, suggesting that high Sle1 levels increase tolerance to oxacillin by promoting cell separation. In contrast, increased oxacillin sensitivity of *sle1* cells appears linked to a synthetical lethal effect on septum synthesis. In conclusion, this study demonstrates that Sle1 is a key factor in resistance to β-lactam antibiotics in the JE2 USA300 model strain, and that PBP2a is required for expression of Sle1 in JE2 cells exposed to oxacillin.

**Importance:** The bacterium *Staphylococcus aureus* is a major cause of human disease, and the global spread of *S. aureus* resistant to β-lactam antibiotics (MRSA) has made treatment increasingly difficult. β-lactams interfere with cross-linking of the bacterial cell wall, however, the killing mechanism of this important class of antibiotics is still not fully understood. Here we provide novel insight into this topic by showing that β-lactam resistance is controlled by the Sle1 cell wall amidase in the fast spreading and highly virulent MRSA USA300 clone. We show that Sle1 high levels accelerate the onset of daughter cells splitting and decrease cell size. Vice versa, oxacillin decreases the Sle1 level, and imposes a cell-separation defect that is antagonized Sle1. The key finding that resistance to β-lactams correlates positively to expression of Sle1 indicates that, in *S. aureus,* the detrimental effects of β-lactam antibiotics are linked to inhibition of daughter cells splitting.

## Introduction

The commensal bacterium *Staphylococcus aureus* that is colonizing the nasal cavity of about one third of the human population is a leading cause of bacterial infections with disease manifestations ranging from superficial skin infections to life-threatening invasive diseases (1). Historically, β-lactam antibiotics have been the agents of choice for the treatment of staphylococcal infections. However, effective treatment of these infections is hampered by the rapid spread of methicillin resistant *S. aureus* (MRSA) that are resistant to virtually all members of the class of β-lactam antibiotics (1, 2). The emergence of community acquired MRSA (CA-MRSA) has dramatically increased the global burden of *S. aureus* infections, and the CA-MRSA clone USA300 is currently the most frequent cause of purulent skin infections in emergency departments in the United States (2, 3). MRSA strains have acquired resistance to β-lactam antibiotics through horizontal acquisition of the *mecA* gene encoding PBP2a, an alternative transpeptidase with low affinity for most β-lactams. Hence, PBP2a is capable of performing the critical cross-linking of peptidoglycan strands when the native penicillin-binding proteins (PBPs) are inhibited by the irreversible binding of β-lactams to the active site (1). Clinically, MRSA isolates exhibit highly variable levels of resistance and specifically USA300 strains exhibit a relatively low level of resistance compared to other MRSA strains (4, 5). The molecular mechanisms underlying the strain dependent resistance to β-lactams remain poorly understood, but the lack of correlation between resistance level and the level of PBP2a expression, suggests that factors other than PBP2a are involved (6–9). One such factor is PBP4 that is required for β-lactam resistance in the CA-MRSA strains MW2 and USA300, but not in the highly resistant hospital-acquired-MRSA strain COL (4, 10).

The highly conserved cytoplasmic ClpXP protease is composed of separately encoded proteolytic subunits (ClpP), and ATPase units (ClpX), where ClpX serves to specifically recognize, unfold, and translocate substrates into the ClpP proteolytic chamber for degradation (11). Interestingly, inactivation of each of the components of the ClpXP protease substantially increased the β-lactam resistance level of the CA-MRSA USA300 model strain JE2 without changing the level of PBP2a, or, the muropeptide profiles of the cell wall, and the mechanism by which ClpXP proteolytic activity modulates β-lactam resistance remained unexplained (5). In *S. aureus,* only a few ClpXP substrates, such as the essential transcriptional regulator, Spx, and the cell wall amidase Sle1, have been identified (12–14). Here we report that the highly increased β-lactam resistance displayed by the USA300 cells lacking ClpXP activity is completely lost upon inactivation of Sle1, suggesting that high Sle1 levels are causing the increased β-lactam resistance of *clpX* or *clpP* mutants. Conversely, inactivation of *sle1* rendered USA300 wild-type cells hyper-sensitive to β-lactam antibiotics. These results are surprising, as the activity of cell wall hydrolases is typically associated with cell lysis following β-lactam treatment, not with promoting survival (15–18). The finding that Sle1 modulates the resistance level of USA300 JE2, prompted us to assess the role of the Sle1 cell wall amidase in S. *aureus* cell division in more detail. Super resolution microscopy revealed that high Sle1 levels accelerate the onset of daughter cell separation starting from the peripheral wall resulting in cells of reduced size. Vice versa, oxacillin imposes a cell-separation defect that is rescued by high Sle1 activity, suggesting that high Sle1 activity enhances tolerance to oxacillin by promoting daughter cell splitting. We further show that expression of Sle1 is correlated to the transpeptidase activity of PBPs, and that PBP2a is required for continued Sle1 expression in cells exposed to oxacillin. Finally, we show the increased oxacillin sensitivity of *sle1* cells seems to be linked to a synergistic lethal effect on septum synthesis.

## Results

### Disruption of the ClpP recognition tripeptide in ClpX confers β-lactam hyper-resistance in USA300

We previously showed that deletion of either the *clpX* or the *clpP* gene resulted in a substantial increase in β-lactam resistance of the clinically important CA-MRSA clone USA300, suggesting that β-lactam resistance can be modulated via pathways depending on the activity of the ClpXP protease (5). In *S. aureus*, ClpP can associate with an alternative substrate recognition factor, ClpC (19), while ClpX independently of ClpP functions as a molecular chaperone (20). To confirm that ClpP and ClpX controls β-lactam resistance via formation of the ClpXP protease, we investigated if β-lactam resistance is increased in cells that retains ClpX chaperone and ClpCP activity but cannot form the ClpXP protease due to a single amino acid substitution in the ClpP recognition IGF motif of ClpX (21). Indeed, introduction of an I_265_E substitution in the IGF tripeptide of ClpX increased the MICs of JE2 against all tested β-lactams confirming that inactivation of the ClpXP protease enhances the β-lactam resistance level of the clinically important CA-MRSA clone USA300 (Table 1). Specifically, expression of the ClpX_I265E_ variant increased the MICs of oxacillin, cefotaxime, and meropenem approximately 8-fold, while causing a minor 2-fold increase in the MICs of imipenem and cefoxitin. We conclude that ClpXP contributes to cellular processes that determine the β-lactam resistance level of JE2.

**Table 1:** β-lactam susceptibility of *sle1*, *clpX_I265E_*, *clpX_I265E_ sle1* and *mecA* mutants in MRSA (JE2) and MSSA (SA564 and 8325-4) wild-type (WT) backgrounds. MIC (μg ml^-1^) is as indicated below.

| Antibiotic | JE2 |  |  |  |  |  | SA564 |  |  | 8325-4 |  |  |
| --- | --- | --- | --- | --- | --- | --- | --- | --- | --- | --- | --- | --- |
|  | WT | *sle1* | *mecA* | *clpX*<sub>I265E</sub> | *clpX*<sub>I265E</sub>, *sle1* | *clpX*<sub>I265E</sub>, *mecA* | WT | *sle1* | *clpX*<sub>I265E</sub> | WT | *sle1* | *clpX*<sub>I265E</sub> |
| Oxacillin | 32-64 | 0.5 | 0.75 | >256 | 0.5 | 1 | 0.5 | 0.5 | 0.75 | 0.38 | 0.25 | 0.5 |
| Imipenem | 0.12 | 0.12 | 0.12 | 0.25 | 0.25 | <0.25 | 0.5 | 0.12 | 0.12 | 0.12 | 0.12 | 0.12 |
| Cefotaxime | 8 | 2-4 | 2 | 64 | 2 | 4 | 4 | 2 | 4 | 1 | 1 | 1 |
| Cefoxitin | 16 | 4 | 4 | 32 | 4 | 4 | 4 | 4 | 4 | 2 | 2 | 4 |
| Cefepime | 4 | 2 | 2 | >32 | 4-8 | 4 | 4 | 4 | 4 | 2 | 1 | 2 |
| Ertapenem | 0.5 | 0.12 | 0.25 | >2 | 0.12 | 0.25 | 0.5 | 0.12 | 0.25 | 0.12 | 0.12 | 0.25 |
| Meropenem | 0.25 | 0.12 | 0.12 | 2 | 0.12 | 0.12 | 0.12 | 0.06 | 0.25 | 0.12 | 0.06 | 0.12 |

### Sle1 is conferring increased β-lactam resistance in JE2 lacking ClpXP activity

In *S. aureus*, the cell wall amidase Sle1 is a substrate of the ClpXP protease, and consequently the cellular levels of Sle1 are elevated in cells lacking ClpXP activity (21). To investigate if the high Sle1 levels play a role in the hyper-resistant phenotype of cells expressing the ClpX_I265E_ variant, we next inactivated *sle1* in JE2 wild-type and *clpX_I265E_* cells and assessed the impact on β-lactam MICs. Interestingly, inactivation of *sle1* not only abrogated the increased resistance of cells lacking ClpXP protease activity, but decreased MICs below the wild-type level (Table 1). Similarly, inactivation of *sle1* in the JE2 wild-type decreased MICs of all β-lactams except imipenem, and rendered JE2 hyper-sensitive to oxacillin, with the oxacillin MIC decreasing from 32 µg ml^-1^ in wild-type cells to 0.5 µg ml^-1^ in *sle1* cells. In fact, inactivation of *sle1* rendered cells as sensitive to oxacillin as did inactivation of *mecA* (Table 1). JE2mecA cells expressing the ClpX_I265E_ variant were as sensitive to β-lactams, as were JE2mecA expressing wild-type ClpX, demonstrating that high Sle1 levels only confer resistance to cells expressing PBP2a. This result is consistent with previous results showing that neither deletion of *clpP* nor deletion of *clpX* alter the MICs of β-lactams in methicillin sensitive *S. aureus* (MSSA) strains (5), and in agreement with this finding, introduction of the *sle1*- and *clpX_I265E_* mutations into the two MSSA strains, SA564 (clinical isolate) and 8325-4 (lab-strain) had only a slight impact on MICS (Table 1): inactivation of *sle1* reduced MICs of most β-lactams about 2 fold, while expression of the ClpX_I265E_ variant did not impact β-lactam MICs in the MSSA strains (Table 1).

Standard MIC-assays prescribe the use of stationary cells and we finally asked, if Sle1 levels also impact the ability of exponentially growing JE2 cells to form colonies in the presence of different concentrations of β-lactams. Consistent with the MIC tests, the spot assay showed that JE2clpX_I265E_ cells were capable of forming colonies in the presence of antibiotic concentrations that inhibited growth of JE2 wild-type cells for all tested β-lactams (Fig. S1). Furthermore, inactivation of *sle1* rendered both wild-type and *clpX_I265E_*cells hyper-sensitive to all tested β-lactams (Fig. S1).

We conclude that in the JE2 MRSA strain, β-Lactam resistance depends on the Sle1 cell wall amidase, and that ClpXP contributes negatively to β-lactam tolerance via degradation of Sle1.

### Population analysis profiles reveal that sle1 cells become homogenously hyper-sensitive to oxacillin

Similar to other CA-MRSA strains, JE2 displays heterogeneity with respect to β-lactam susceptibility, meaning that the majority of cells exhibit a low level of antibiotic resistance, while a minority of cells is highly resistant (1, 5). In order to determine if inactivation of Sle1 or ClpXP alters the hetero-resistant phenotype of the JE2 strain, a <u>p</u>opulation <u>a</u>nalysis <u>p</u>rofile (PAP) was performed. In the PAP analysis, we chose to focus on oxacillin and cefoxitin, as these two compounds represent β-lactams whose MICs were highly and marginally effected, respectively, by expression of the ClpX_I265E_ variant. As expected, the PAP analysis resulted in a typical heterogeneous profile for the JE2 wild-type strain with the majority of cells being killed by low concentrations of either oxacillin or cefoxitin, while a small subpopulation was capable of growing at much higher concentrations of antibiotics (Fig. 1). Expression of the ClpX_I265E_ variant not only increased the fraction of JE2 cells able to grow in the presence of medium high levels (16-32 µg ml^-1^) of antibiotics by 4 logs, but also enabled the most resistant subpopulation to grow at even higher concentrations of antibiotics, Fig. 1. On the contrary, inactivation of *sle1* transformed both the JE2 wild-type and JE2clpX_I265E_ into a homogeneously sensitive strain with all cells in the population being inhibited by very low concentrations of antibiotics, Fig. 1.

**Fig 1.**
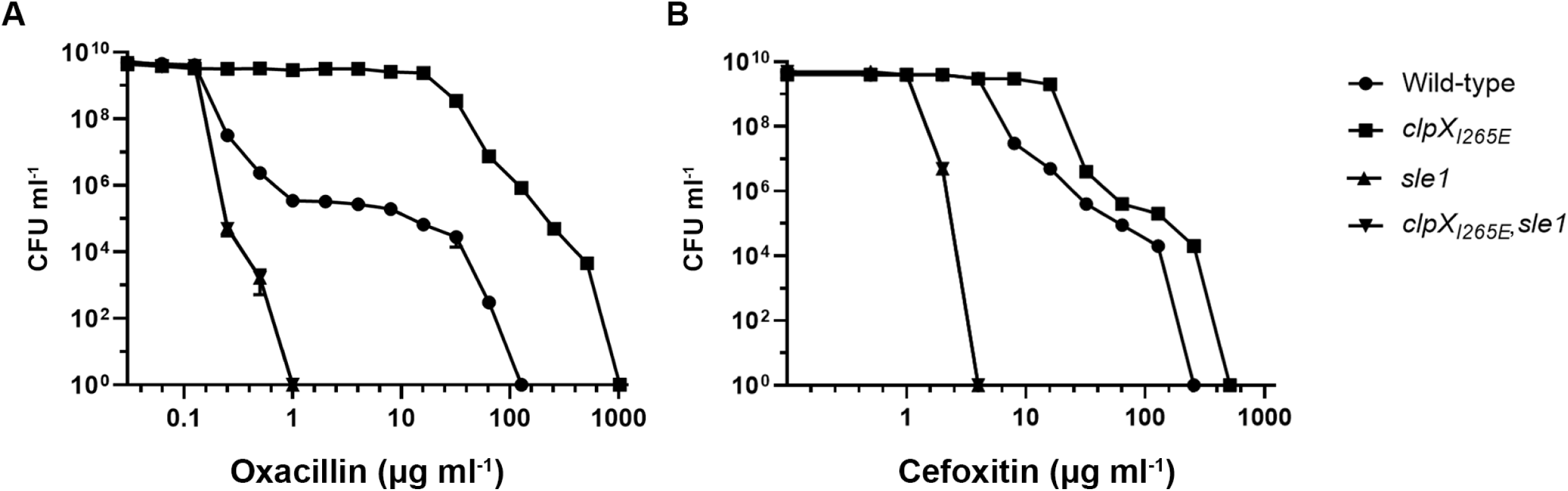
Population analyses profiles show that inactivation of *sle1* renders the JE2 wild-type and JE2clpX_I265E_ homogenously sensitive to oxacillin and cefoxitin. CFU ml^-1^ for JE2 wild type, JE2clpX_I265E_, JE2sle1 and JE2clpX_I265E_,sle1 were determined after plating on increasing concentrations of oxacillin (A), and cefoxitin (B), as indicated. Representative data from three individual experiments are shown.

### Inactivation of Sle1 delays the onset of daughter cell splitting, while high Sle1 levels accelerate the onset of daughter cell separation

Based on the finding that deletion of *sle1* induced formation of cell clusters it was proposed that Sle1 is involved in separation of *S. aureus* daughter cells (14). The interesting finding that Sle1 activity impacts resistance to β-lactam antibiotics prompted us to assess the role of Sle1 in *S. aureus* cell division in more detail using Super-Resolution Structured Illumination Microscopy (SR-SIM). Prior to SR-SIM, cells were stained with the membrane stain, Nile Red, and the cell wall stain fluorescent wheat germ agglutinin (WGA-488) that is too big to penetrate into cells and therefore only labels cell wall exposed to the exterior milieu during the staining period (22, 23). To visualize regions of new peptidoglycan insertion, cells were additionally stained with the blue fluorescent D-amino acid, hydroxycoumarin-amino-D-alanine (HADA). To investigate the impact of Sle1 on the *S. aureus* cell cycle, we first assigned wild-type and mutant cells to different phases based on the state of septum ingrowth (22): newly separated daughter cells that have not initiated septum formation were assigned to phase 1, cells in the process of synthesizing division septa were assigned to phase 2, while cells displaying a closed septum were assigned to phase 3. As depicted in Fig. 2A inactivation of *sle1* significantly increased the fraction of phase 3 cells (P=0.04), while conversely, the fraction of phase 3 cells was significantly reduced in *clpX_I265E_* cells (P = 0.004), however, only if cells express Sle1. As the percentage of cells observed in each growth phase should be proportional to the fraction of the cell cycle spent in that stage, this finding indicates that separation of fully divided daughter cells is delayed in the absence of Sle1, while *clpX_I265E_*cells spend less time in phase 3. In support here off, splitting of the HADA-stained septal wall was only observed in 7% of *sle1* cells, as compared to 31% of wild-type cells, and 69% of cells expressing the ClpX_I265E_ variant – Fig. 3A-B, and D. In conclusion, daughter cell separation is delayed in the absence of Sle1, while high levels of Sle1 seem to accelerate the onset of *S. aureus* daughter cell splitting.

**Fig 2.**
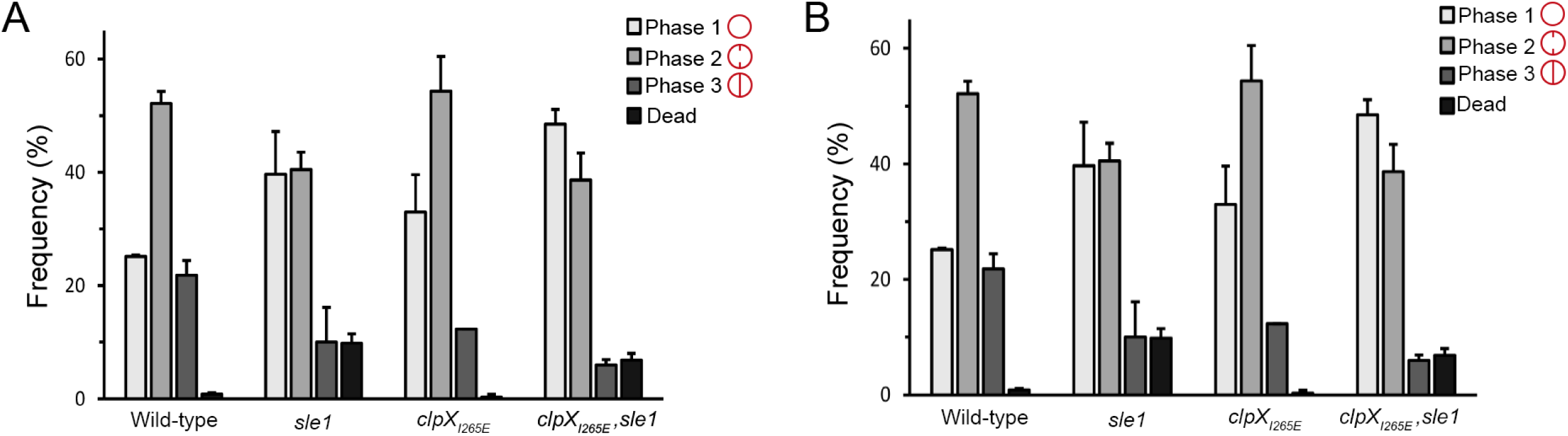
Sle1 and oxacillin impact the *S. aureus* cell cycle. JE2 wild-type, JE2clpX_I265E_, JE2sle1, and JE2clpX_I265E,_ sle1 were grown exponentially at 37°C in the absence (A) or presence of 0.05 μg ml^-1^ oxacillin (B); cells were then stained with membrane dye Nile Red (red) before imaging by SR-SIM. To assess the effect of the mutations on progression of the growth cycle, 300 cells (from each of two biological replicates) were scored according to the stage of septum ingrowth: no septum (phase 1), incomplete septum (phase 2), or non-separated cells with complete septum (phase 3).

**Fig 3.**
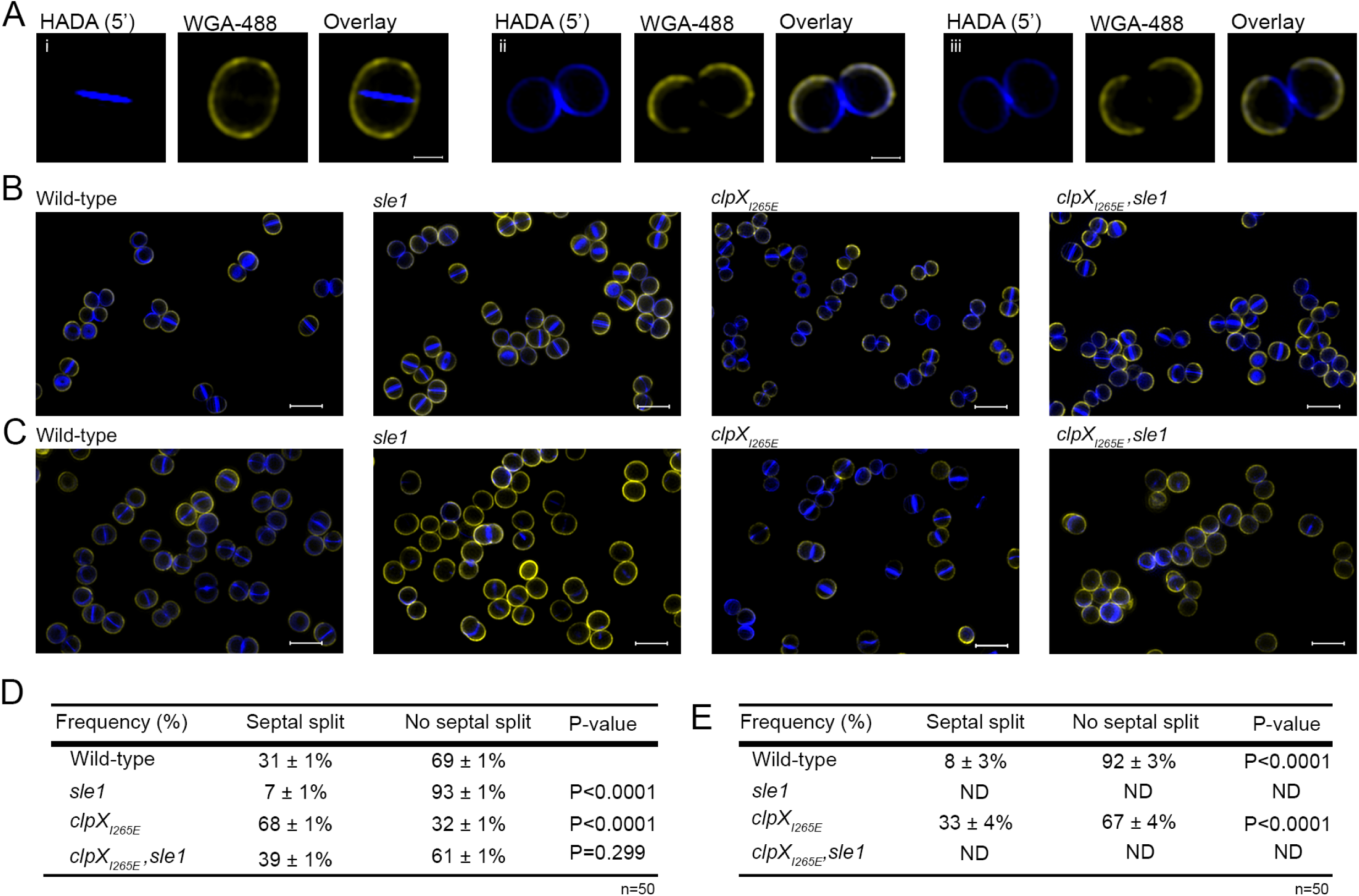
High Sle1 levels accelerate splitting of *S. aureus* daughter cells, while inactivation of Sle1 and oxacillin delays splitting of fully divided daughter cells. JE2 wild type, JE2sle1, JE2clpX_I265E_ and JE2clpX_I265E_, sle1 were grown exponentially at 37°C in the absence (A-B) or presence of 0.05 µg ml^-1^ oxacillin (C); cells were then stained for 5 minutes with WGA-488 (green) that stains the old peptidoglycan, and HADA (blue), that is incorporated into newly synthesized peptidoglycan. Cells were imaging by SR-SIM. The dual peptidoglycan staining allowed us to directly identify daughter cells pairs that have separated following WGA/HADA staining, as these daughter cells display a characteristic labeling pattern with the “old” peripheral cell wall stained in green, while the “novel” peripheral cell wall of septal origin is stained in blue (22, 23). A) displays example images of cells that have completed septum synthesis during the labeling period and have either i) not separated following labeling or ii-iii) initiated daughter cell separation following labeling (shown from two different orientations). Images shown in B and C is displayed with the same intensity of WGA and HADA signal, respectively, with the exception of the WGA signal in JE2clpX_I265E_ cells that has been increased due to the otherwise weak signal in this mutant – both in the absence or presence of oxacillin. D-E) 50 cells that completed septum formation during the labeling period was scored according to splitting of the newly synthesized HADA or no splitting. D) P-values were obtained by Chi-square test against the wild-type. E) P-values were obtained by Chi-square test against the sample grown in the absence of oxacillin.

### Sle1 controls cell size

Characterization of the *S. aureus* cell cycle revealed that *S. aureus* cells are capable of elongating, and that elongation mainly occurs in phase 1 and phase 3 (22, 23). After establishing that the level of Sle1 impacts the time cells spend in phase 3, we determined if Sle1 activity impacts the cell size. Indeed, the estimation of cell size demonstrated that *clpX_I265E_* cells are significantly smaller than wild-type cells (P < 0.0001), whereas *clpX_I265E_, sle1* cells are of similar size as wild-type cells (Fig. 4A). Inactivation of *sle1* in wild-type cells resulted in cells being slightly, but significantly, larger than wild-type cells (P < 0.0001), Fig. 4A. We conclude that high levels of the Sle1 cell wall amidase leads to a decrease in cell size, while inactivation of Sle1 increases the cell size.

**Fig 4.**
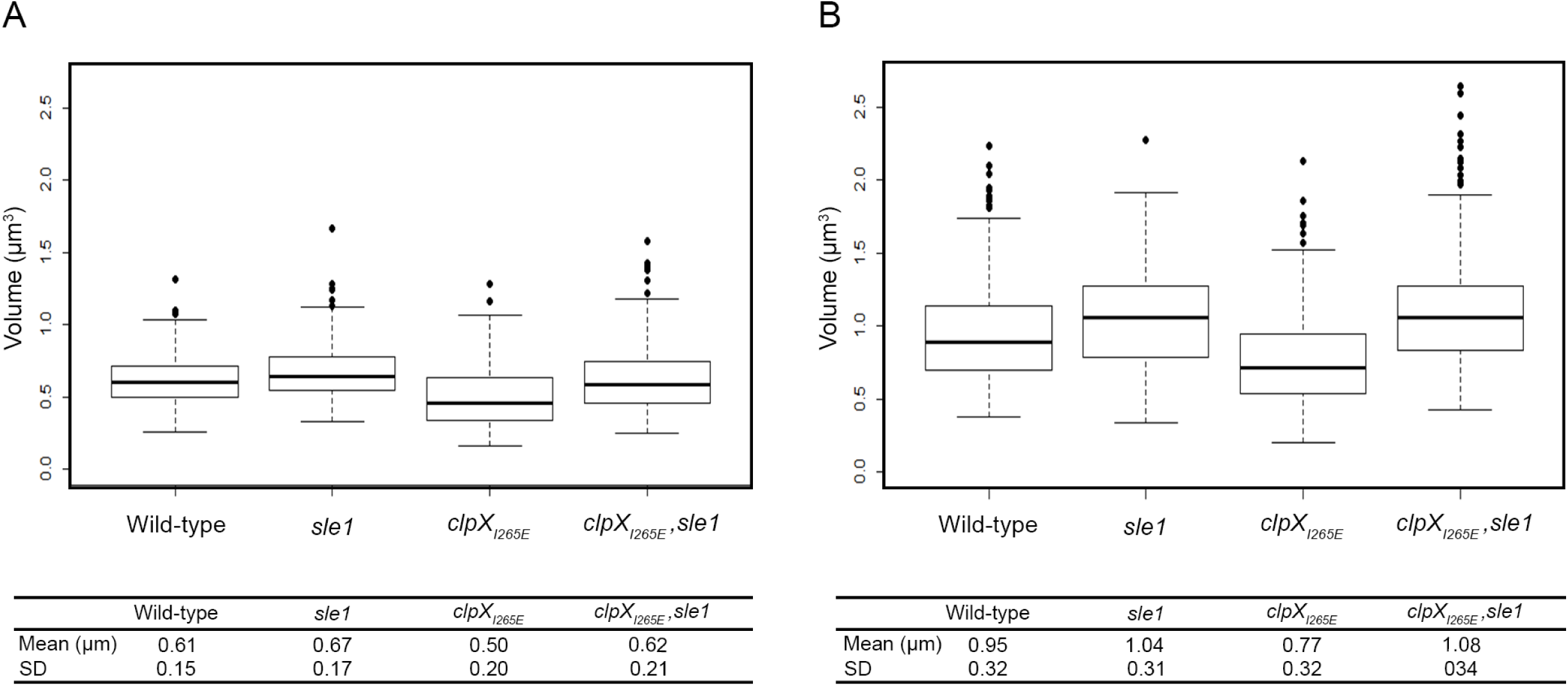
High Sle1 activity reduces the cell size, while oxacillin increases cell size. JE2 wild type, JE2clpX_I265E_, JE2sle1, JE2clpX_I265E_, sle1 were grown exponentially at 37°C in the absence (A) or presence of 0.05 µg ml^-1^ oxacillin (B); cells were then stained with membrane dye Nile Red (red) before imaging by SR-SIM. Cell volume off 300 cells representing 100 cells from each of the three growth phases was determined by fitting an ellipse to the border limits of the membrane. Graph represents data from two biological replicates.

### Scanning electron microscopy indicates that Sle1 contributes to the formation of perforations in the peripheral septal ring prior to popping

At the time of cell separation, *S. aureus* daughter cells are connected only at the edge of the septum by a peripheral ring (23, 24). Resolution of this peripheral wall by mechanical crack propagation results in ultrafast splitting of daughter cells in a process designated “popping” (23). Scanning electron microscopy (SEM) have revealed that popping is preceded by the presence of perforation holes around the bacterial circumference coincident with the outer edge of the division septum, and it was speculated that autolysins are involved in formation of these holes (23). To examine if Sle1 is the autolysin responsible for creating perforation holes along the septal ring, SEM was used to image the cell surface of wild-type and *clpX_I265E_*cells, as well as the surface of the corresponding *sle1*-negative strains (Fig. 5). In agreement with published data, small holes are visible at mid-cells in a small fraction of wild-type cells (Fig. 5). Typically, perforation holes were apparent on ellipsoid cells displaying a slight invagination at mid-cell, supporting that these cells are in the process of dividing. In support of Sle1 being involved in generating these perforation holes, inactivation of *sle1* in both wild-type and *clpX_I265E_*cells rendered the cell wall at mid-cell appear more smooth and less perforated (Fig. 5). On the other hand, the fraction of cells displaying cracks at mid-cell appeared substantially increased in cells expressing the ClpX_I265E_ variant, and most *clpX_I265E_*cells captured in SEM images are in different stages of cell-separation, Fig. 5. These phenotypes of the *clpX_I265E_* cells disappeared upon inactivation of Sle1. We conclude that the fraction of cells displaying perforations at mid-cells correlates to the level of Sle1 expression indicating that the Sle1 cell wall amidase contributes to degradation of the cell wall in the peripheral ring of the division septum prior to popping.

**Fig 5.**
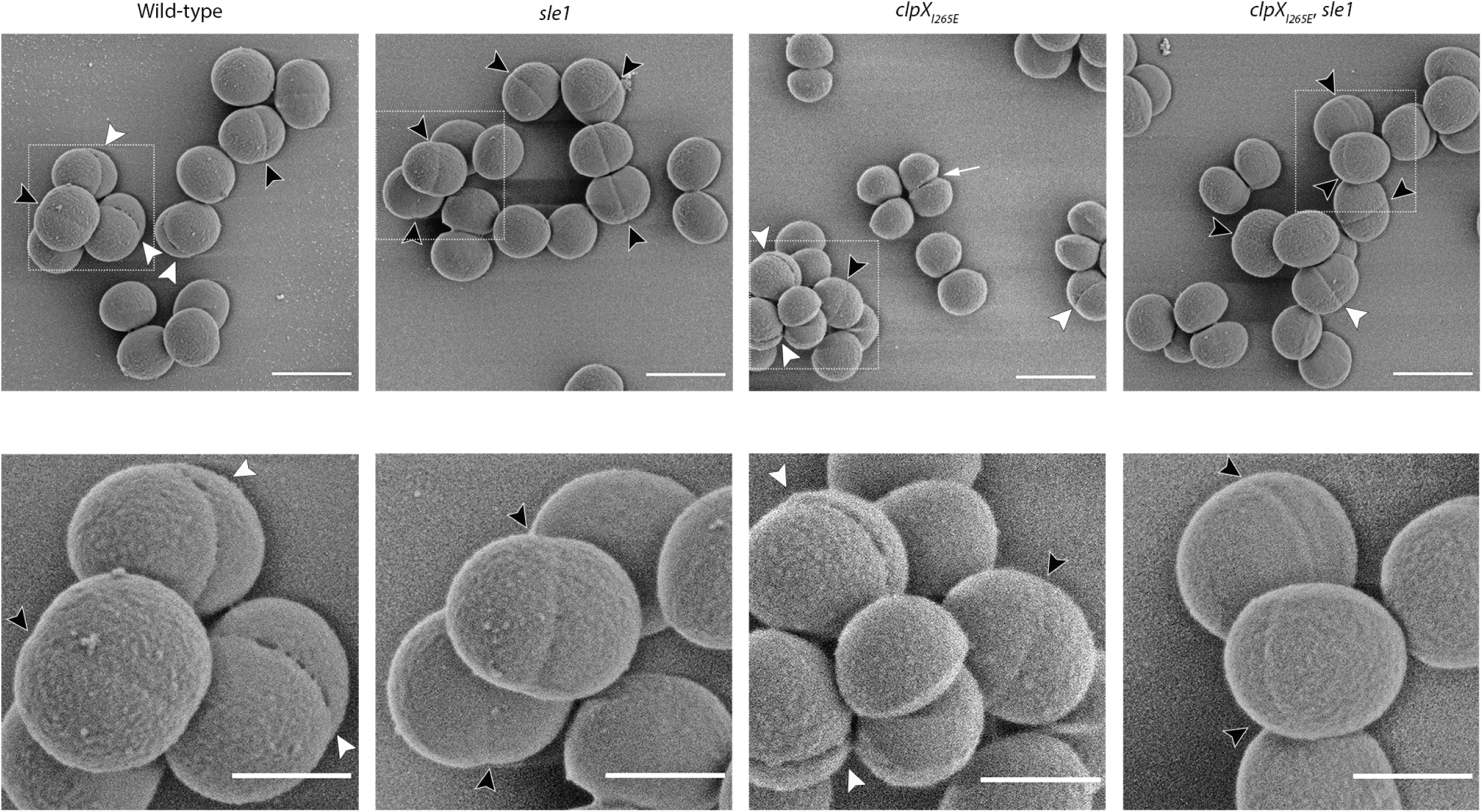
Perforations in the peripheral septal wall correlate with Sle1 levels. SEM images of JE2 wild-type, *sle1*, *clpX_I265E_*or *clpX_I265E_*, *sle1* grown in TSB to mid-exponential phase at 37°C. The images show invaginations (black arrowhead) and perforations (white arrowhead) at mid-cell along the septal ring. The larger cracks in *clpX_I265E_*, is indicated with a white arrow. Scale bars, 1 μm (overview) and 0.5 μm (zoom).

### Oxacillin treatment impedes separation of daughter cells and interferes with septum formation in cells devoid of Sle1

We now sought an answer to why β-lactam resistance correlates to expression of Sle1 in the JE2 background. Based on the finding that Sle1 is required for fast separation of *S. aureus* daughter cells, we first hypothesized that β-lactams impose a cell-separation defect that, while being rescued by high Sle1 activity, is lethal to cells lacking Sle1 activity. To test this hypothesis JE2 wild-type, *sle1* and *clpX_I265E_*cells were exposed to 1.25 µg ml^-1^ oxacillin (1/4 MIC of cells devoid of Sle1 activity) for 20 min before cells were stained with Nile Red, WGA, and HADA and imaged by SR-SIM. Consistent with the idea that oxacillin impedes splitting of daughter cells, the fraction of wild-type cells displaying splitting of the HADA stained septum was significantly reduced from 31% to less than 10% (P< 0.0001) upon exposure to oxacillin (Fig. 3C and E). Likewise, the fraction of wild-type cells divided by a closed septum (phase 3 cells), and the size of wild-type cells increased significantly following oxacillin exposure (Fig. 2 and Fig. 4, respectively). Moreover, high Sle1 levels seems to antagonize the cell separation defect conferred by oxacillin, as significantly more *clpX_I265E_*than wild-type were capable of splitting in the presence of oxacillin (Fig. 2 and Fig. 3). Taken together these data support that oxacillin confers a cell separation defect that is rescued by high Sle1 activity. On the other hand, oxacillin was expected to exacerbate the cell separation defects of cells devoid of Sle1, however, oxacillin rather diminished the fraction of *sle1* cells in phase 3 (Fig. 2), and very few oxacillin treated *sle1* cells displayed a closed HADA-stained septum (Fig. 3). In fact, the HADA signal was weak, or even absent in the majority of *sle1* cells exposed to oxacillin (Fig. 3; Fig. S2). Strikingly, the ability of cells to incorporate HADA in the presence of oxacillin correlated with Sle1 expression, as oxacillin also reduced the HADA signal in wild-type cells, while the intensity of the HADA signal in JE2clpX_I265E_ was similar +/- oxacillin exposure, and did not change, even if cells were treated with higher oxacillin concentrations (Fig. 3; Fig. S2). Therefore, oxacillin seems also to impose a septum synthesis defect that is exacerbated in the absence of Sle1. In support here off, SR-SIM images revealed abnormal septal ingrowths in 75% of *sle1* cells exposed to oxacillin (Fig. 6; Fig. S2). To study the morphological changes induced by oxacillin in cells lacking Sle1 in more detail, cells exposed to oxacillin for 20 min were additionally imaged with transmission electron microscopy (TEM). Both SR-SIM and TEM confirmed severe abnormalities in septal ingrowths in oxacillin treated *sle1* cells, with septa being devoid of the electron-dense septal mid-zone previously designated “the splitting line” (25) (Fig. 6C, i-iv; Fig. S2, and Fig. S3), septa protruding asymmetrically inwards (Fig. 6C, iii-iv), and displaying a characteristic “curvy” morphology (Fig. 6C, ii-iv). Both TEM and SR-SIM images also revealed that exposure to oxacillin resulted in lysis of a small fraction of *sle1* cells, and that lyzed *sle1* cells were typically observed in daughter cell pairs, where the lyzed cell is attached to a living cell (Fig. 6C, iv). Similar, but less severe changes in the septum morphology were observed in wild-type exposed to the same concentration of oxacillin (Fig. 6A; Fig. S2, and Fig. S3). In summary, we found that while high levels of Sle1 seem to enhance tolerance to oxacillin by antagonizing an oxacillin induced cell separation defect, the increased oxacillin sensitivity of *sle1* cells seems to be linked to a synthetical lethal effect on septum synthesis.

**Fig 6:**
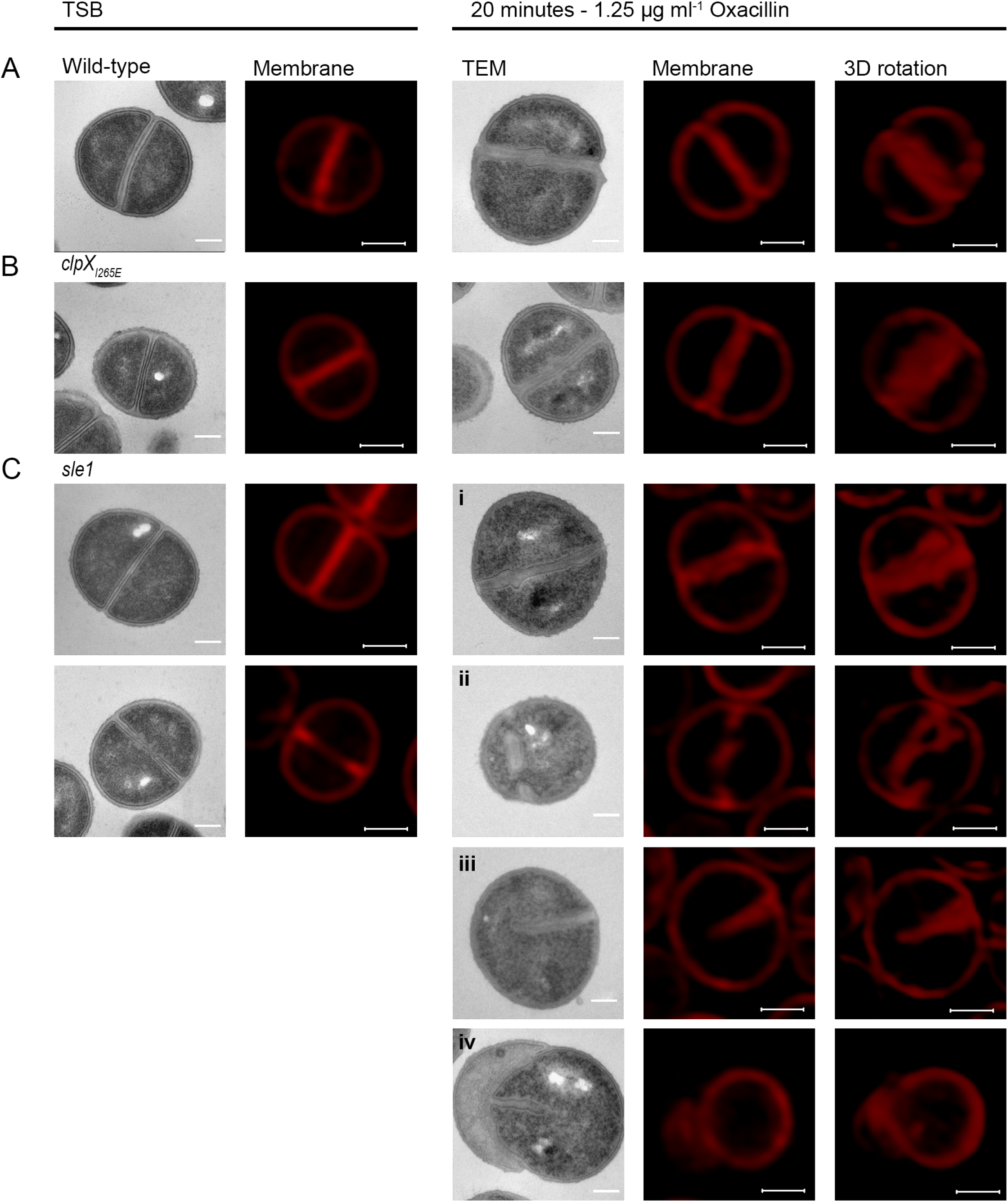
TEM and SR-SIM reveal severe septal abnormalities in *sle1* cells exposed to oxacillin. TEM and SR-SIM images of JE2 wild-type (A), *clpX_I265E_* (B) or *sle1* (C) grown to mid-exponential phase at 37°C in the absence (left panels) or presence of 1.25 µg ml^-1^ oxacillin for 20 minutes (right panels) as indicated. Images show the characteristic changes induced by oxacillin determined from at least two biological replicates. (A,B) In JE2 wild-type and JE2clpX_I265E_ cells, oxacillin treatment leads to septal abnormalities including curving and thickening of the septa and blurring of the electron-dense mid-zone. (C) In JE2sle1 cells, the effect of oxacillin is exaggerated; i-iv) septa are devoid of the electron-dense mid-zone, ii) septa appearing not to be attached to the septal wall, but 3D rotation reveal a non-circular septal pattern, iii) septa protruding asymmetrically inwards, and iv) lysis in daughter cell pairs where the lyzed cell is attached to a living cell. Scale bar, TEM; 0.2 μm, SR-SIM; 0.5 μm.

### Expression of Sle1 is reduced in cells exposed to oxacillin

The two major cell wall hydrolases involved in *S. aureus* daughter cell splitting are Sle1 and Atl (14,26,27). To examine if oxacillin interferes with splitting of JE2 daughter cells by reducing expression of these enzymes, murein hydrolase activity was determined in cell wall extracts derived from cells grown in the absence or presence of oxacillin using standard zymography. The resulting murein hydrolase profiles showed multiple bands that were apparently AtlA-related, since they disappear in extracts from an Δ*atl* mutant (Fig. 7A; Fig. S4A). Sle1 is a 32 kDa protein comprised of an N terminal cell wall binding domain and a C terminal catalytic domain with N-acetyl muramyl-l-alanine amidase activity (14). In the zymographs, the activity of the Sle1 autolysin is also clearly visible, and as expected cell wall extracts derived from JE2clpX_I265E_ cells displayed higher Sle1 activity than wild-type cells both in the absence or presence of oxacillin (Fig. 7A).

**Fig 7.**
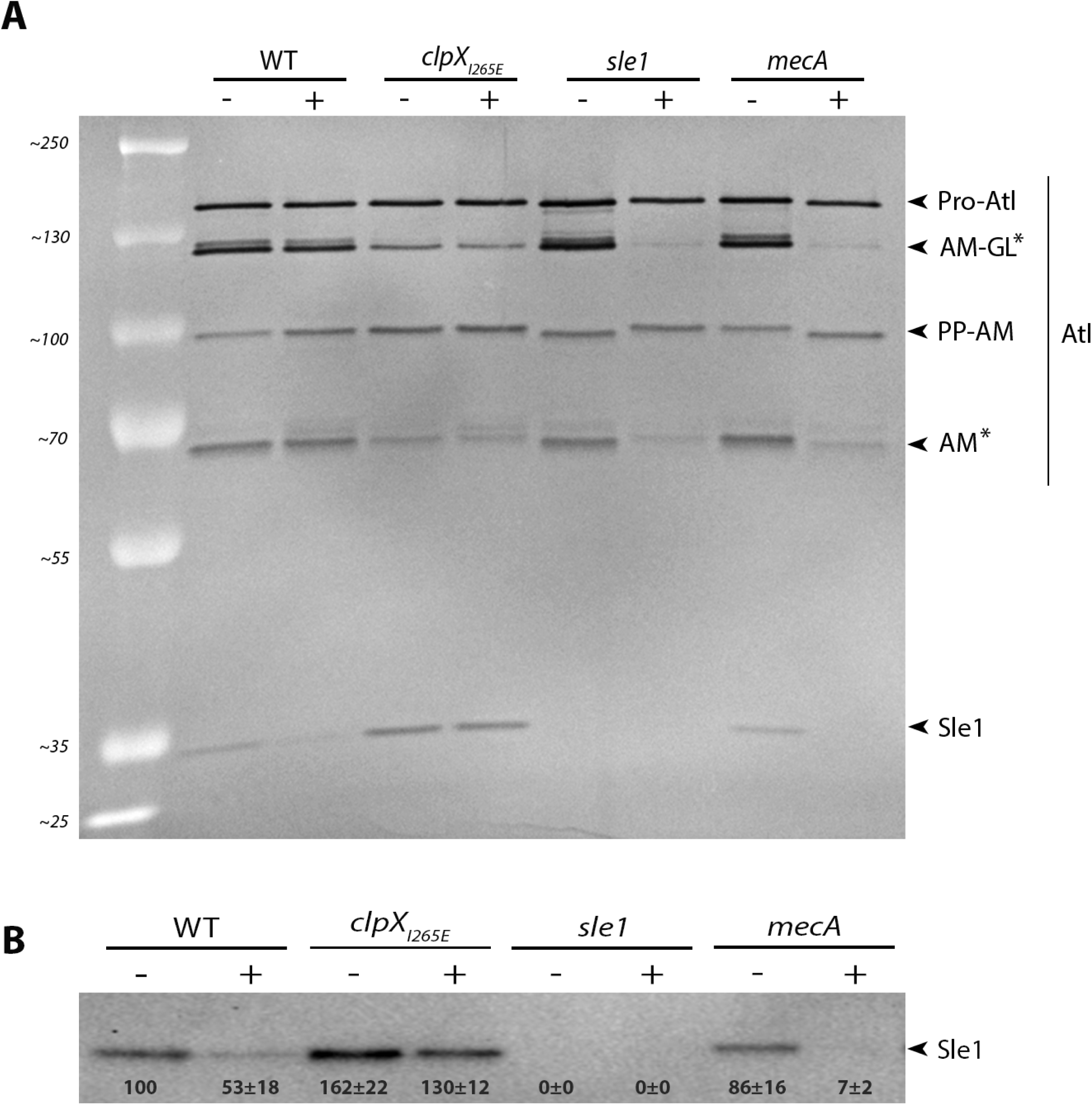
Oxacillin reduces expression of Sle1, and PBP2a is required for expression of Sle1 in JE2 cells exposed to oxacillin. (A) Zymogram assay. JE2 and its derived strains were grown to exponential phase in the absence (-) or presence (+) of 8 µg ml^-1^ oxacillin as indicated, and equal amounts of cell wall-associated proteins were loaded onto an SDS polyacrylamide gel containing heat-killed S. aureus JE2 cells. One representive gel of two biological independent experiments is shown as an inverted image of the strained gel. The positions of the molecular mass standards are indicated (in kilodaltons) on the left. Asterisks indicates Atl bands that are diminished in oxacillin exposed JE2 cells lacking Sle1 or PBP2A. (B) Sle1 levels in cell wall extracts were additionally determined by Western blot analysis in three biological replicates. Densitometry analysis was performed using Fiji. Obtained values were normalized to values obtained for the wild-type and are displayed below the corresponding bands.

Interestingly, only the intensity of the Sle1 band was diminished in cell wall extracts derived from JE2 cells exposed to oxacillin (Fig. 7A). These findings indicate that Sle1 expression is down-regulated, or, alternatively that export of Sle1 to the cell wall is reduced in JE2 exposed to oxacillin. To distinguish between these two possibilities, we additionally determined the Sle1 level in cell wall extracts and in whole cells by Western blotting (Fig. 7B; Fig. S4B). In agreement with previous findings, both the non-exported and the exported form of Sle1 accumulate in JE2 cells expressing the ClpX_I265E_ variant [21]. In wild-type cells, however, the non-exported Sle1 was neither detected in cells grown in the absence, nor in cells grown in the presence of oxacillin (Fig. S4B), and a similar 2-fold reduction in the Sle1 level were observed in cell wall extracts and in extracts from whole cells (Fig. 7B; Fig. S4B). Hence, oxacillin seems to reduce expression, not export of Sle1.

### Sle1 levels are reduced 10-fold in JE2mecA cells exposed to oxacillin

The finding that exposure to oxacillin reduced levels of Sle1 in JE2 cells shows that expression of Sle1 is positively coupled to the trasnpeptidase activity of PBPs. Oxacillin blocks the transpeptidase domain of native PBPs and, therefore, we predicted that expression of Sle1 depends on the transpeptidase activity of PBP2a in oxacillin-treated JE2 cells. To test this hypothesis, we determined the levels of Sle1 in JE2mecA cells grown in the absence or presence of oxacillin. Interestingly, this analysis revealed that, whilst the intensities of Sle1 bands were similar in extracts from wild-type and *mecA* cells grown in the absence of oxacillin, Sle1 was barely detectable in *mecA* cells exposed to oxacillin (Fig. 7). Taken together, these results lend support to the idea that Sle1 expression is coupled to activity of the transpeptidase domain of PBPs, and that Sle1 expression becomes dependent on PBP2a activity in JE2 cells exposed to oxacillin.

Finally we noted that while oxacillin did not impact intensities of the different Atl bands in JE2 wild-type cells, the intensities of two Atl bands were diminshed in oxacillin exposed JE2 cells not expressing Sle1 or PBP2A (Fig. 7A, asterisks). The bifunctional Atl murein hydrolase is produced as a precursor protein (Pro-Atl) that undergoes proteolytic cleavage to yield two catalytically active proteins, an amidase (AM) and a glucosaminidase (GL) (27). The 113 kDA band repesents an intermediary cleavage product, while the 62 kDa band reflects activity of the fully cleaved amidase (28). Therefore, PBP2a and Sle1 may additionally have a role in Atl processing in oxacillin exposed JE2 cells.

## Discussion

β-lactam antibiotics are the most frequently prescribed antibiotics world-wide, however, the mechanism by which the binding of β-lactams to their PBP targets causes death and lysis of bacteria is not completely understood (1, 29,30). In at least some bacteria, the killing mechanism of β–lactams involves unsynchronized activation of peptidoglycan hydrolases (15–18). Contradictive to this model, we here show that the activity of the Sle1 cell wall amidase is crucial for PBP2a mediated resistance to β-lactams in the JE2 CA-MRSA model strain, and that elevated levels of Sle1 confers increased resistance to β-lactams, however, only if JE2 expresses PBP2a. The key finding that resistance to β-lactams correlates positively to expression of Sle1 indicates that, in *S. aureus,* the detrimental effects of β-lactam antibiotics are linked to inhibition, rather than to activation, of peptidoglycan hydrolase activity.

Sle1 was proposed to function in *S. aureus* cell division (14), and with the recent advances in SR-SIM we now examined the role of Sle1 in this fundamental process in more detail using cells either lacking or over-producing Sle1. To divide, *S*. *aureus* builds a septal cross wall generating two hemispherical daughter cells that, at the time of cell separation, are connected only at the peripheral ring forming the outer edge of the septum, (23, 24). Resolution of this peripheral wall involves mechanical crack propagation, but the contribution of cell wall hydrolases to the ultra-fast popping of *S. aureus* daughter cells remains poorly described (23). We demonstrate that high levels of Sle1 accelerate the onset of daughter cell separation starting from the peripheral wall indicating that Sle1 contributes to the timely degradation of the outer edge of the septal wall. Sle1 is, however, not required for resolution of the outer septal wall, as separation of daughter cells is delayed not inhibited in cells lacking Sle1. Previously, cryo-electron microscopy revealed that, at the beginning of septation, the peripheral ring is thicker that other parts of the outer wall, and it was proposed that this extra cell wall material serve to protect the peripheral wall from degradation by the cell wall hydrolases functioning in presplitting of the septal cross-walls (24). Interestingly, the TEM and SR-SIM pictures presented here suggest that Sle1 degrades the peripheral wall from the exterior not from the interior. Taken together, our data support that high Sle1 levels promote daughter cell splitting, hence, indicating that the detrimental effect of β-lactam antibiotics is linked to impaired daughter cell separation.

We, similarly to others, observed that oxacillin delays cell separation (18,31,32), and based on the characteristic “hour-glass” morphology observed for wild-type cells exposed to oxacillin (indicating that splitting from the cell periphery has taken place, Fig. S3), we speculate that oxacillin primarily interferes with splitting of the interior septal cross-walls. Interestingly, the electron dense line that vanishes in oxacillin exposed cells was previously described as tubular packets enclosing autolytic enzymes that upon completion of the cross wall are released to facilitate cell separation (18, 25). Next, we showed that oxacillin reduces expression of Sle1 in JE2 cells, suggesting that oxacillin impairs daughter cell splitting by down-regulating Sle1 expression.

Interestingly, the Tomasz lab has convincingly shown that transcription of genes encoding cell wall hydrolases is tightly linked to activities of the PBPs, and that inhibition of PBP activity, either by genetic depletion or by treatment with β-lactam antibiotics, reduces transcription of a number of cell wall hydrolase genes including *sle1* (33–35). In these studies, the β-lactam induced transcriptional repression of cell wall hydrolase genes was proposed to be a defence mechanism protecting cells with perturbed cell wall synthesis from the destructive forces of cell wall hyrolases. Based on our paradoxical finding that high levels of Sle1 increase resistance of *S. aureus* to β-lactam antibiotics, we instead hypothesize that the shut-down of transcription of *sle1* and other cell wall hydrolase genes in *S. aureus* cells exposed to β-lactam antibiotics is part of the mechanism that eventually end up killing the cells. Instead, we speculate that PBP-mediated transpeptidation, the last step in peptidoglycan synthesis, is involved in signaling that peptidoglycan synthesis is complete, and that it is time to activate expression of *sle1* and other cell wall hydrolases (Model depicted in Fig. 8). According to this model, the non-native PBP2a has an important role in activating expression of Sle1 in cells, where the transpeptidase activity of native PBPs is inhibited by the irreversible binding of oxacillin to the active site (Fig. 8). Consistent with this model, we here show that while PBP2a is required for expression of Sle1 in JE2 cells exposed to oxacillin, deletion of *mecA* does not impact Sle1 levels in JE2 cells grown in the absence of oxacillin. The observation that Sle1 levels are reduced 2-fold upon exposure to oxacillin indicates that the non-native PBP2a is less efficient in promoting expression of Sle1 (Fig. 8). In *S. aureus*, at least 13 genes encode known or putative peptidoglycan hydrolases (36). Transcription of many of these genes respond to PBP activity in a strain-dependent manner (33–35), hence, we speculate that the strain dependent resistance level, conferred by *mecA*, is linked to the ability of PBP2a to co-ordinate expression of cell-wall hydrolases with septum synthesis.

**Fig 8.**
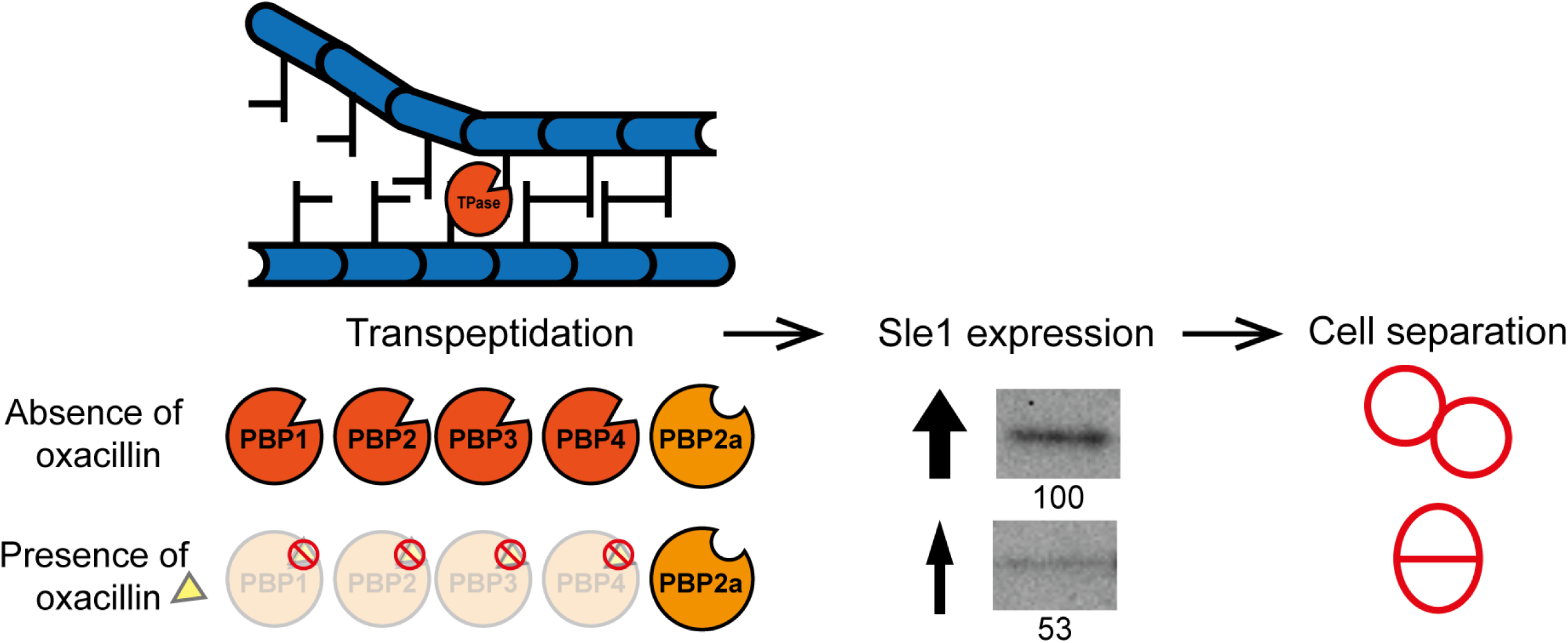
Sle1 in S. aureus daughter cell splitting, model. Model of delayed daughter cell splitting in the absence of Sle1 and accelerated splitting in cells with high Sle1 levels. In wild-type cells, progression of the cell cycle goes from spherical cells with no septum (phase 1) to formation (phase 2) and closing/ completion of the septum (phase 3). Following this, resolution of the peripheral wall and subsequent rapid popping of daughter cells takes place. In the absence of Sle1, cells stay longer in phase 3 and some daughter cells initiate formation of a new septum before separation of daughter cells has occurred. However, in *clpX_I265E_* that has high levels of Sle1, the onset of daughter cell splitting is accelerated and cells spend less time in phase 3. Illustration made by Esben Thalsø-Madsen.

Finally, we found that inactivation of *sle1* is synthetically lethal with sub-mic concentrations oxacillin in JE2. At present we cannot explain this finding, but the severe septum defects conferred by oxacillin in cells devoid of Sle1 activty indicate that Sle1 and the transpeptidase activity of PBPs function synergistically to coordinate septum formation with daughter cell separation. Consistent with this finding, Matias and Beveridge (24) proposed that cell wall autolysins are required for synchronized growth of the septum. Intriguingly, a recent paper suggests that another *S. aureus* cell wall amidase, LytH, is involved in controlling the spatial distribution of peptidoglycan synthases to ensure that cell expansion is coordinated with cell division (37). Hence, the activities of cell wall hydrolases and PBPs may be tightly linked to balance peptidoglycan synthesis with autolytic degradation, and the binding of β-lactam antibiotics PBPs may perturb this delicate balance.

## Methods

### Bacterial strains and growth conditions

Bacterial strains used in this study are listed in table 2. *S. aureus* JE2, SA564 and 8325-4 were used as wild-type strains. Strains were cultured in 20 ml tryptic soy broth (TSB; Oxoid) with shaking (170 rpm) at 37°C or on tryptic soy agar (TSA; Oxoid) at 37°C. In all experiments, bacterial strains were freshly streaked from the frozen stocks on TSA and incubated overnight at 37°C. From these plates, TSB cultures were inoculated to an OD_600_ of 0.05 or below and optical densities (OD) measured at 600 nm.

**Table 2:** Bacterial strains used in the present study

| Strain | Description | Source |
| --- | --- | --- |
| **JE2** | CA-MRSA strain USA300 LAC cured of plasmids | (38) |
| **JE2 *sle1*** | *sle1*::$\Phi\text{N}\Sigma$ transduced from NE1688 (Fey *et al.*, 2013) into JE2 wild-type | This study |
| **JE2 *clpX*<sub>I265E</sub>** | *clpX*<sub>I265E</sub> variant from 8325-4 *clpX*<sub>I265E</sub> transduced into JE2 wild-type | This study |
| **JE2 *clpX*<sub>I265E</sub>, *sle1*** | *sle1*::$\Phi\text{N}\Sigma$ transduced from NE1688 (Fey *et al.*, 2013) into JE2 *clpX*<sub>I265E</sub>. *ermB* | This study |
| **JE2 *mecA*** | *mecA*::$\Phi\text{N}\Sigma$ transduced from NE1868 (Fey *et al.*, 2013) into JE2 wild-type. *ermB* | This study |
| **JE2 *clpX*<sub>I265E</sub>, *mecA*** | *mecA*::$\Phi\text{N}\Sigma$ transduced from NE1868 (Fey *et al.*, 2013) into JE2 *clpX*<sub>I265E</sub>. *ermB* | This study |
| **SA564** | Low-passage clinical isolate | (39) |
| **SA564 *sle1*** | *sle1*::$\Phi\text{N}\Sigma$ transduced from NE1688 (Fey *et al.*, 2013) into SA564 wild-type. *ermB* | This study |
| **SA564 *clpX*<sub>I265E</sub>** | *clpX*<sub>I265E</sub> variant from 8325-4 *clpX*<sub>I265E</sub> transduced into SA564 wild-type | This study |
| **8325-4** | Widely used wild-type strain cured of all prophages | (40) |
| **8325-4 *sle1*** | *clpX*<sub>I265E</sub> variant from 8325-4 *clpX*<sub>I265E</sub> transduced into 8325-4 wild-type. *ermB* | This study |
| **8325-4 *clpX*<sub>I265E</sub>** | 8325-4 expressing a *ClpX*<sub>I265E</sub> variant from the native *clpX* locus | (21) |

### Construction of strains

Sle1 and PBP2a were inactivated by introducing *sle1*::ΦΝΣ from NE1688 or *mecA*::ΦΝΣ from NE1868, respectively, (38) using phage 85 and selecting for resistance to erythromycin. In order to construct a JE2clpX_I268E_, sle1 double mutant, JE2 chromosomal *clpX* was first replaced with an untagged version of *clpX_I265E_* (substituting the ATT codon with GAA*ref) by phage 85 mediated transduction using 8325-4clpX_I265E_ (21) as the donor. In order to select for JE2 cells that had incorporated the marker-less *clpX_I265E_* variant, we took advantage of the finding that expression of the ClpX_I265E_ variant increases the resistance of JE2 to oxacillin by plating tranductants at 50 µg ml^-1^ oxacillin. Introduction of the *clpX_I265E_* variant was subsequently confirmed using a primer pair designed to distinguish wild-type *clpX* from *clpX_I265E_* (3’ end is complementary to the mutated GAA codon, sequence underlined in primer: clpX_I265E__F (5’-CGT CTT GGT GAA AAA GTT <u>GAA</u>) and clpX_R (5’-CCG TGG CTA GCA TGT TTA AAT TCA ATG AAG A). To ensure that the high oxacillin concentration had not introduced additional mutations in the genome of the JE2clpX_I265E_ candidate, the genomes of the JE2 wild-type and the JE2clpX_I265E_ candidate were sequenced by Illumina sequencing on a NextSeq instrument at the Danish National Reference Laboratories for Resistance Surveillance (SSI, Copenhagen, Denmark). All sequence analysis was performed in CLC Genomics Workbench Software, version 12.0 (https://www.qiagenbioinformatics.com). The sequencing reads from JE2 wild-type and the JE2clpX_I265E_ candidate were mapped to USA300 FPR3757 chromosomal genome sequence (GenBank accession number NC_007793.1). Genomic variations were identified using the “Basic Variant Detection” tool in CLC. This analysis confirmed the introduction of the GAA codon in JE2clpX_I265E_ candidate, and identified three additional SNPs between the JE2 wild-type sequence and the sequence of the JE2clpX_I265E_ candidate: one SNP was found in a non-coding region close to *clpX*, while the two others SNPs mapped in the *hemA* gene that is located one gene downstream of *clpX*. The *hemA* sequence in the JE2_clpXI265E_ candidate is identical to the *hemA* sequence in 8325-4, suggesting that all identified SNPs originate from 8325-4clpX_I265E_ that was used as the donor in the transduction.

### Susceptibility testing

Susceptibility testing was performed by the Danish National Reference Laboratories for Resistance Surveillance (SSI, Copenhagen, Denmark) using Etest® (bioMérieux) to determine the MICs of oxacillin and including *S. aureus* strain ATCC43300 as a reference strain. The Sensititre™ Vizion™ broth microdilution system (Thermo Fisher Scientific) was used to determine the MICs of all other antibiotics using *S. aureus* strain ATCC29213 as a reference strain.

### Population analysis profiles (PAP)

Population analysis profiles were determined by plating appropriate dilutions of an overnight *S. aureus* culture on TSA plates containing increasing concentrations of oxacillin or cefoxitin (Sigma). Plates were incubated at 37°C for 48 h and the number of colonies was determined and plotted against antibiotic concentration as described previously (Sieradzki *et al.* 1998).

### SR-SIM analysis

#### Imaging and sample preparation

For SR-SIM analysis, cells were imaged with an Elyra PS.1 microscope (Zeiss) using a Plan-Apochromat 63x/1.4 oil DIC M27 objective and a Pco.edge 5.5 camera. Images were acquired with five grid rotations and reconstructed using ZEN software (black edition, 2012, version 8.1.0.484) based on a structured illumination algorithm, using synthetic, channel specific optical transfer functions and noise filter settings ranging from -6 to -8. Laser specifications can be found in Table 3. Prior to imaging, cultures of *S. aureus* were grown at 37°C for four generations before dividing the cultures into two; one grown in the absence of oxacillin and the other supplemented with 1.25 µg ml^-1^ oxacillin. Cultures where grown for 20 minutes before sample preparation. Cells were stained at 37°C for 5 min with the membrane dye Nile Red, the cell wall dye WGA-488 and the fluorescent D-amino acid HADA (Table 3). Samples were washed twice in PBS, placed on an agarose pad (1.2% in PBS), and visualized by SR-SIM as described above. SR-SIM analysis was performed at the Core Facility of Integrated Microscopy (CFIM).

**Table 3:** Staining and laser specifications used for SR-SIM

| Staining | Concentration | Target | Laser | Laser Type | Laser power | Beam splitter | Grating |
| --- | --- | --- | --- | --- | --- | --- | --- |
| **Nile Red** | 5 $\mu\text{g ml}^{-1}$ | Membrane | 561 nm | HR Diode – 100mW | 5 % | BP 570-650 + LP 750 | 34 $\mu\text{m}$ |
| **HADA** | 250 $\mu\text{M}$ | Old PG | 405 nm | HR Diode – 50mW | 20 % | BP 420-480 + LP 750 | 23 $\mu\text{m}$ |
| **WGA-488** | 1 $\mu\text{g ml}^{-1}$ | New PG | 488 nm | HR Diode – 100mW | 5 % | BP 495-575 + LP 750 | 28 $\mu\text{m}$ |

#### Analysis of the cell cycle

To address progression of the cell cycle, 300 cells were scored according to the stage of septum ingrowth: no septum (phase 1), incomplete septum (phase 2), or non-separated cells with complete septum (phase 3). Scoring was based on the Nile Red staining. Dead cells were scored as collapsed cells with no HADA incorporation. To enumerate the fraction of phase 2 cells displaying symmetric septal ingrowth versus abnormal ingrowths, 100 Nile-Red stained cells displaying septal ingrowths were evaluated in each of two biological replicates. To quantify daughter cell splitting, 50 cells that had completed septum formation during the 5 minutes of HADA labeling were scored based on whether they displayed no septal splitting as depicted in Fig. 3A, i or septal splitting as depicted in figure Fig. 3A, ii-iii. This analysis was performed for two biological replicates.

#### Estimating cell volume (size)

The volume of 300 cells representing 100 cells from each of the three different growth phases was determined (two biological replicates). An equal number of cells from each phase was used in order to avoid bias in average volume due to a shift in the phase distribution. Volume was determined as described in (Monteiro *et al*., 2015). Briefly, an ellipse was fitted to the border limits of the membrane to acquire measurements of the minor and major axis. The cell shape was assumed to be that of a prolate spheroid and the volume was estimated using the equation V = 4/3πab^2^; where a and b correspond to the major and minor axes, respectively. Ellipse fitting and measurements were performed using Fiji (http://fiji.sc).

### Statistical analysis

All statistical analysis were performed using R statistical software. Student’s t-test was used to assess significant differences in cell volume. The Chi-squared test of independence was used to determine if there was a significant relationship between the proportion of cells assigned to each of the three phases or relevant phenotypes under the tested condition (number of cells in the relevant phase or phenotype/the total number of cells). A value P < 0.05 was considered significant.

### Electron microscopy

#### Scanning electron microscopy (SEM)

Strains were grown in TSB at 37°C as specified with an initial starting OD of 0.02. Exponentially growing cells were collected and placed on ice for 5 min prior to centrifugation (13400 rpm; 1 min). Cell pellets were resuspended in fixation solution (2% glutaraldehyde in 0.05 M sodium phosphate buffer, pH 7.4) and deposited on the glass discs at 4°C for a minimum of 24 h. The specimens were subsequently washed three times in 0.15 M sodium phosphate buffer (pH 7.4) and post-fixed in 1% OsO_4_ in 0.12 M sodium cacodylate buffer (pH 7.4) for two hours. Following a rinse in distilled water, the specimens were dehydrated to 100% ethanol according to standard procedures and critical point dried (Balzers CPD 030) with CO2. The specimens were subsequently mounted on stubs using double adhesive carbon tape (Ted Pella) as an adhesive and sputter coated with 6 nm gold (Leica ACE 200). SEM observations were performed using a FEI Quanta 3D scanning electron microscope operated at an accelerating voltage of 2 kV.

#### Transmission electron microscopy (TEM)

Strains were grown in TSB at 37°C as specified above, but with an initial OD_600_ of 0.02 and at OD_600_ = 0.2 the culture was divided in two; one without oxacillin and one supplemented with 1.25 µg ml^-1^ oxacillin. The cultures were incubated at 37°C for 20 minutes. Cells were collected by centrifugation (8000 g; 5 min) and suspended in fixation solution as above and incubated overnight at 4°C. The fixed cells were further embedded in agarose, rinsed three times in 0.15 M sodium phosphate buffer (pH 7.2), and subsequently post-fixed in 1% OsO_4_ with 0.05 M K_3_Fe(CN)_6_ in 0.12 M sodium phosphate buffer (pH 7.2) for two hours. The specimens were dehydrated in a graded series of ethanol, transferred to propylene oxide and embedded in Epon according to standard procedures. Sections, approximately 60 nm thick, were cut with an Ultracut 7 (Leica, Wienna, Austria) and collected on copper grids with Formvar supporting membranes, stained with uranyl acetate and lead citrate. Specimens examined with a Philips CM 100 Transmission EM (Philips, Eindhoven, The Netherlands), operated at an accelerating voltage of 80 kV. Digital images were recorded with an Olympus Veleta digital slow scan 2.048 x 2.048 CCD camera and the ITEM software package.

All SEM and TEM processing and microscopy of fixed cells were performed at the CFIM.

### Western blotting

*S. aureus* cultures were inoculated in TSB at 37°C and grown for four generations. Cultures were divided into two; one without oxacillin and one supplemented with 8 µg ml^-1^ oxacillin. The cultures were further incubated at 37°C for 60 minutes and cells were harvested at OD_600_ of 1-2. Cell wall-associated proteins were extracted by resuspending pellet in 4% SDS (normalized to OD 1 ml^-1^) and incubated for 45 min at room temperature (37°C) with gentle shaking as described previously (41). For immunoblotting, samples were loaded on NuPAGE 10% Bis-Tris gels (Invitrogen™) using MOPS-Buffer (Invitrogen™). After separation, proteins were blotted onto a polyvinylidene difluoride (PVDF) membrane using the Invitrolon™ PVDF Filter Paper Sandwich (0.45 µm pore size, Invitrogen™). Membranes were pre-blocked with Human IgG to avoid a Protein A signal. The Sle1 protein was detected using rabbit-raised antibodies against staphylococal Sle1 (14). Bound antibody was detected with the WesternBreeze Chemiluminescent Anti-Rabbit kit. Densitometry analysis for three biological replicates was performed using the ImageJ “Gel Analysis tool”, where the background from the gel was removed individually for each band

### Zymographic analysis

Bacteriolytic enzyme profiles were obtained using a 10% SDS-PAGE with embedded heat-killed (125°C for 15 min) *S. aureus* JE2 wild-type cells as substrate. Autolytic enzyme-extracts were prepared by growing bacterial strains as described for western blotting. 10 ml of culture was withdrawn and washed twice with 1 volume of ice-cold cold 0.9% NaCl (FK). Cell wall-associated proteins were extracted by resuspending pellet in 1 ml of 4% SDS (normalized to OD 1 ml^-1^) and incubating for 45 min at room temperature (37°C) with gentle shaking. Cells were precipitated (8000 rpm; 5 min) and the supernatant used as a source of enzymes. Following electrophoresis, the gel was washed with ionized water for 15 min three times and subsequently incubated for 20-24 hours in renaturing buffer (50 mM Tris-HCl (pH 7.5), 0.1% Triton X-100, 10 mM CaCl2, 10 mM MgCl2) at 37°C with gentle agitation. The gel was rinsed in ionized water, stained (0.4% methylene blue, 0.01% KOH, 22% EtOH) for 1 min, and destained with ionized water for 1 h with gentle agitation prior to photography.

## Acknowledgments

We greatly acknowledge Professor Simon Foster (University of Sheffield) for the generous gift of FDAA’s, the Nebraska Transposon Mutant Library (NTML) for providing strains, and Motoyuki Sugai (Hiroshima University), for providing Sle1 antibodies. Finally, we would like to thank the staff at the Core Facility for Integrated Microscopy (University of Copenhagen) for their enthusiastic assistance in doing SEM, TEM and SR-SIM.

---

**S1 Fig.**
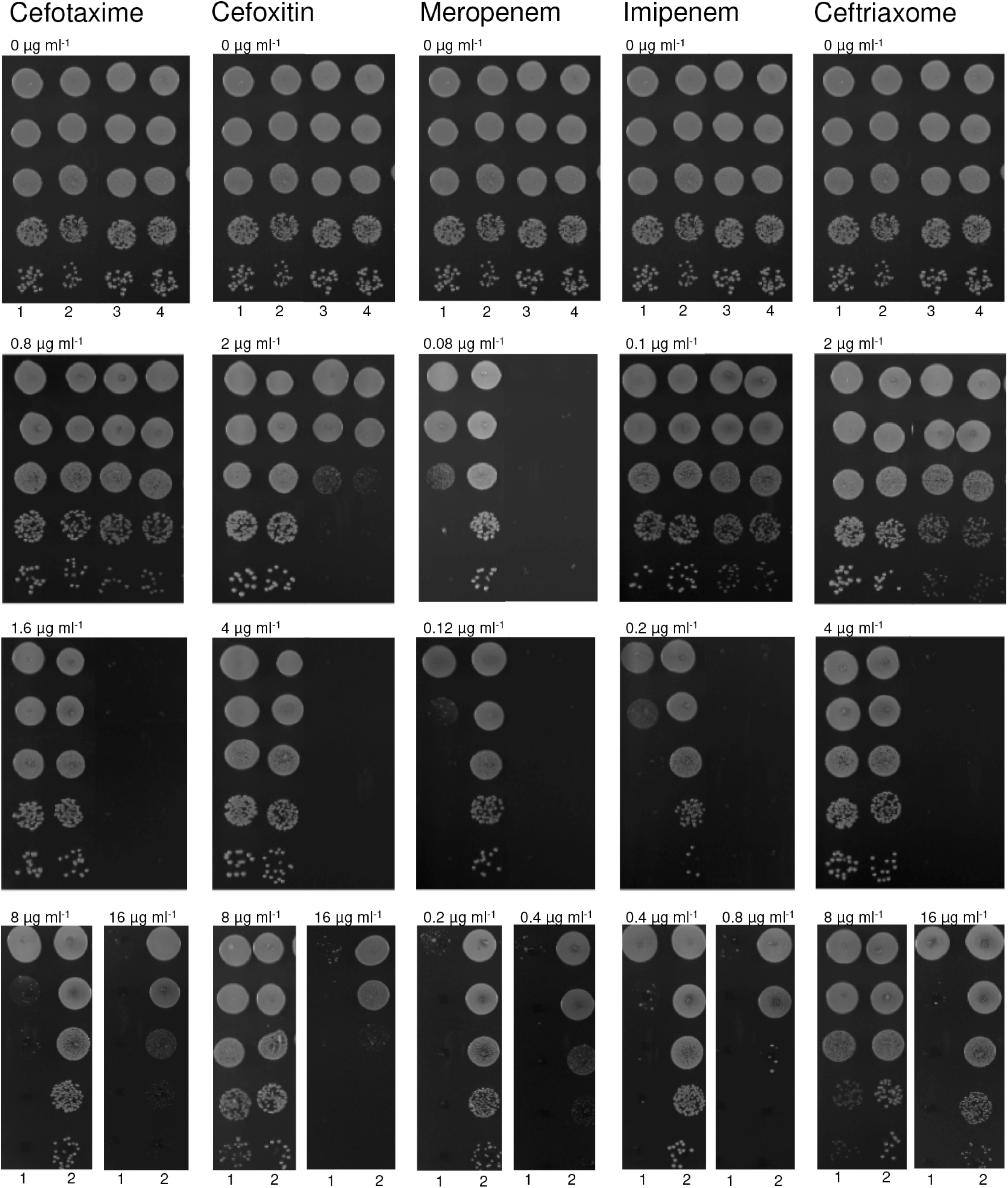
β-lactam susceptibility in JE2 is correlated with Sle1 levels. *S. aureus* JE2 wild-type (1), *sle1* (2), *clpX_I265E_* (3) and *clpX_I265E_ sle1* (4) were grown exponentially in TSB at 37°C. At OD_600_ = 0.5, cultures were diluted 10^1^, 10^2^, 10^3^ and 10^4^- fold. 10 μl of 10^0^ and each dilution were spotted on TSA plates with or without β-lactams as indicated.

**Fig S2:**
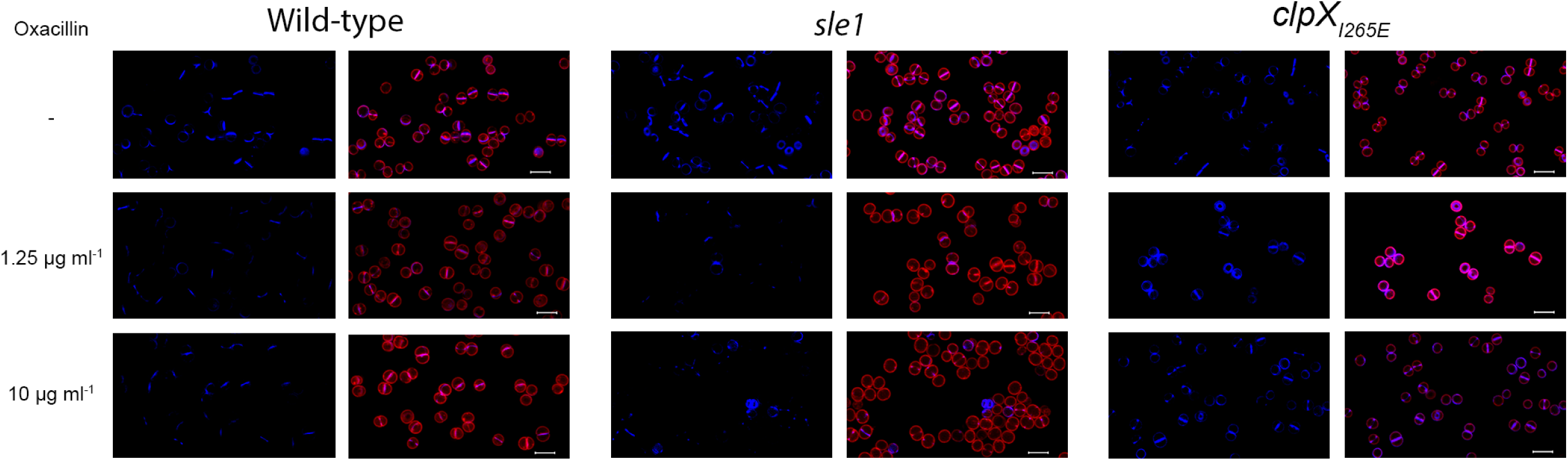
Oxacillin interferes with peptidoglycan synthesis in cells devoid of Sle1. SR-SIM images of JE2 wild-type, *clpX_I265E_,* or *sle1* grown to mid-exponential phase at 37°C in the absence or presence of 1.25 µg ml^-1^ or 10 µg ml^-1^ oxacillin for 20 minutes as indicated. White asterisk indicates examples of cells with no active peptidoglycan synthesis (no HADA signal) and green asterisk indicates examples lyzed cells.

**Fig. S3.**
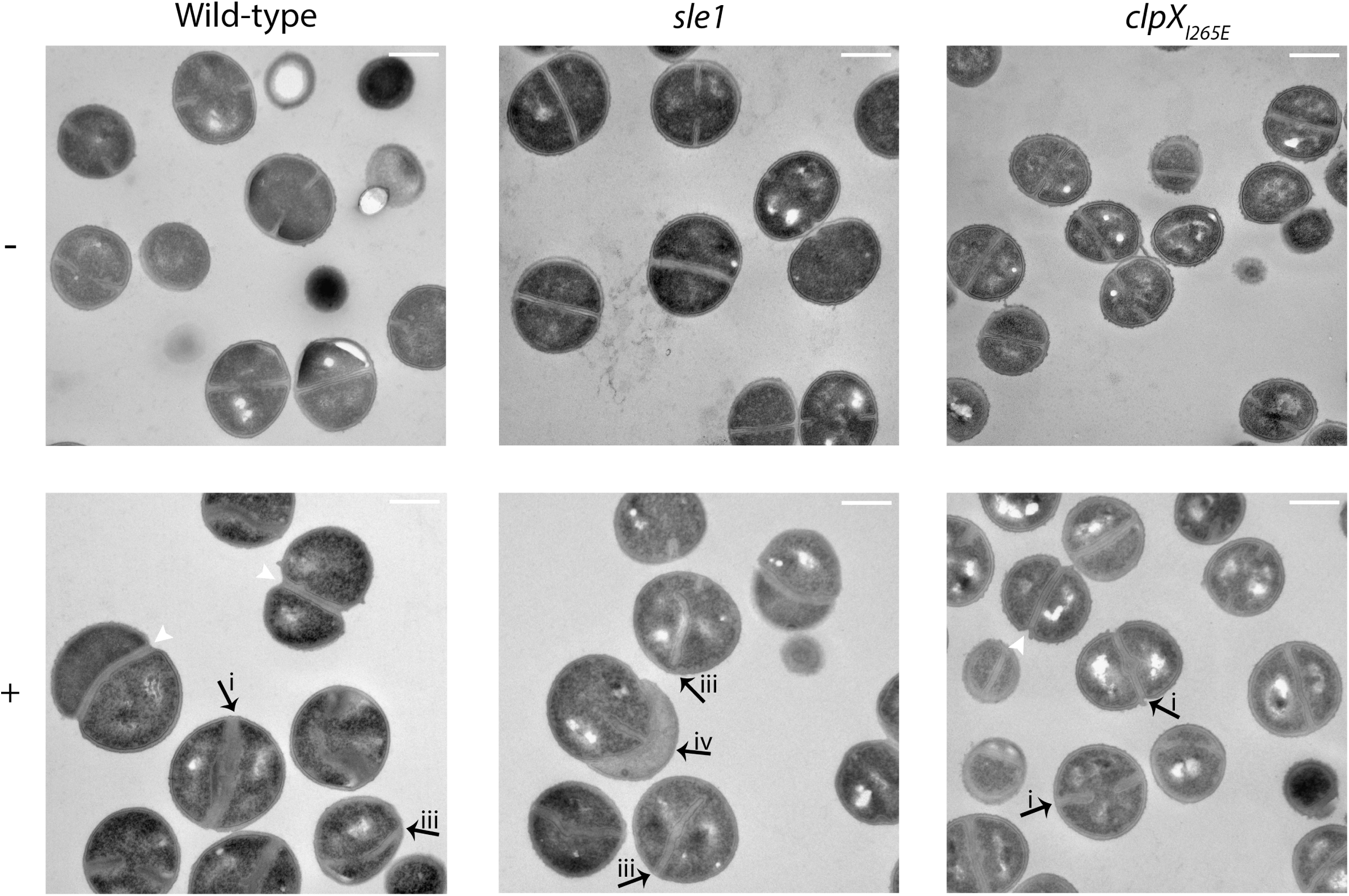
TEM images showing morphological changes in cells exposed to oxacillin. TEM images of JE2 wild-type, *sle1* or *clpX_I265E_*grown in TSB to mid-exponential phase at 37°C in the absence (-) or presence of 1.25 µg ml^-1^ oxacillin (+). Images show several characteristic morphologies of β-lactam treated wild-type and mutant cells also described in Fig. 6 indicated with black arrows (i-iv). These include i-iv) septa devoid of the electron-dense mid-zone; ii) septa appearing not to be attached to the septal wall; iii) septa protruding asymmetrically inwards often only from one side and iv) lysis of cells occurring in daughter cell pairs where the lyzed cell is still attached to a living cell. The “hour-glass” figure is indicated with a white arrowhead. The scale bar corresponds to 0.5 μm.

**Fig S4.**
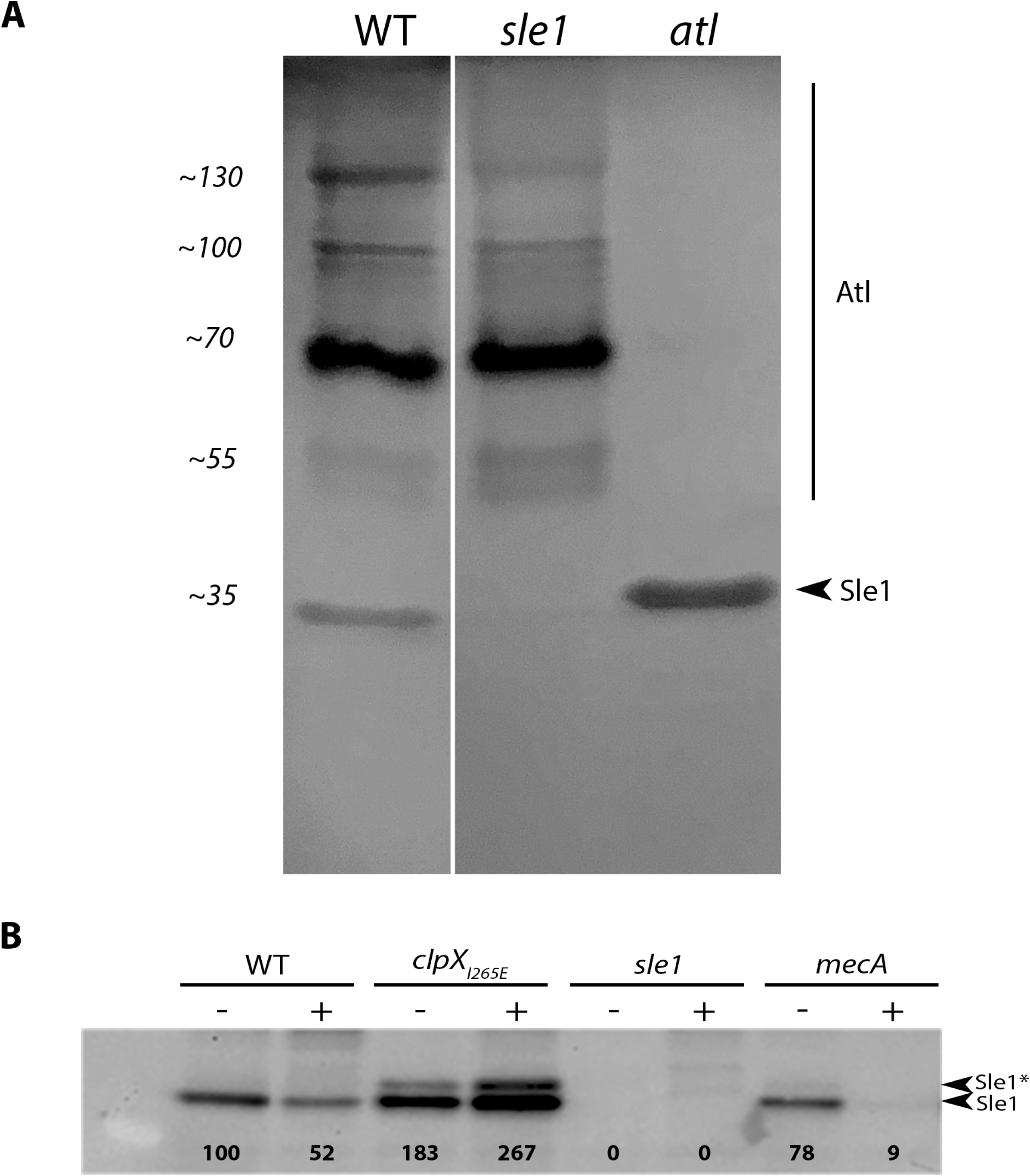
Sle1 does not accumulate intracellularly in cells exposed to oxacillin. (A) Zymography demonstrating that the activity of Atl is visible in multiple bands reflecting that the bifunctional Atl murein hydrolase is produced as a precursor protein (Pro-Atl) that is sequentially cleaved to generate several intermediates (B) Sle1 levels in whole cell extracts from JE2 wild-type, *clpX_I265E_*, *sle1* and *mecA* grown at 37°C in the absence (-) or presence (+) of 8 μg/ml oxacillin, were determined by western blotting using a Sle1 specific antibody. Densitometry analysis was performed using ImageJ, and values were normalized wild-type values and is displayed below the bands. In contrast to cell wall fractions (Fig. 7), the Sle1 antibody recognizes two bands of similar sizes in the whole cell extract that both disappears in the *sle1* mutant. Based on these observations we speculate that they represent Sle1 with (Sle1*) and Sle1 without a signal peptide attached.

